# Sexual selection and the evolution of male and female cognition: a test using experimental evolution in seed beetles

**DOI:** 10.1101/514711

**Authors:** Julian Baur, Jean d’Amour, David Berger

## Abstract

“The mating mind hypothesis”, originally aimed at explaining human cognition, holds that the socio-sexual environment shapes cognitive abilities among animals. Similarly, general sexual selection theory predicts that mate competition should benefit individuals carrying “good genes” with beneficial pleiotropic effects on general cognitive ability. However, few experimental studies have evaluated these related hypotheses due to difficulties of performing direct tests in most taxa. Here we harnessed the empirical potential of the seed beetle study system to investigate the role of sexual selection and mating system in the evolution of cognition. We evolved replicate lines of beetle under enforced monogamy (eliminating sexual selection) or polygamy for 35 generations and then challenged them to locate and discriminate among mating partners (male assays) or host seeds (female assays). To assess learning, the same beetles performed the task in three consecutive rounds. All lines learned the task, improving both within and between trails. Moreover, polygamous males outperformed monogamous males. However, there were no differences in the rate of learning between males of the two regimes, and polygamous females showed no improvement in host search, and even signs of reduced learning. Hence, while sexual selection was a potent factor that increased cognitive performance in mate search, it did not lead to the general increase in cognitive abilities expected under the “mating mind” hypothesis or general “good genes” theory. Our results highlight sexually antagonistic (balancing) selection as a potential force maintaining genetic variation in cognitive traits.

## Background

Cognitive traits allow for behavioural plasticity which can fundamentally change evolutionary dynamics and the mode of, and limits to, adaptation^1–4^. Cognitive abilities also vary widely among animal taxa and there are many hypotheses aimed at explaining this interspecific variation. Most evolutionary explanations typically emphasize the importance of trade-offs and species ecology in shaping cognition^5–10^. One such hypothesis is the idea that the social system of a species is particularly important in shaping cognitive abilities^11^. Indeed, the complex social structure of human societies has been suggested as a main driver of our species’ intelligence^12^. This idea has also been expanded and popularized to include the socio-sexual environment, advocating the view that sexual selection and competition over mating partners has been an important factor contributing to human cognition, which only later allowed the successful colonization of new environments and unprecedented cultural innovations of our species (“*The mating mind hypothesis*”)^13,14^.

The arena for socio-sexual interactions and associated cognitive decision making appear somewhat different in humans compared to other animals. However, it is perhaps only from an anthropocentric perspective that these differences can be seen as larger than those between any other two animals with different mating systems^15^, suggesting that the mating mind hypothesis could be generalized to explain variation among non-human taxa^16^. Indeed, the idea that sexual selection requires cognitive abilities has been widely explored (e.g. ^5,14,16–18^). However, comparative evidence for a direct link between mating system variation and animal cognition is mixed^16,17,19^. For example, in primates monogamy rather than polygamy is associated with larger brain size^20^, providing evidence against the hypothesis that sexual selection leads to increased cognition but not excluding that complex social structure is important, given that maintaining monogamous pair-bonds may be cognitively demanding^5,11^.

In both primates^6^ and birds^21^, environmental complexity is more strongly associated with brain size than the social system of the species. Moreover, in bats, males of species where females are promiscuous tend to have smaller brains but larger testes compared to species where females exhibit mate fidelity^22^, suggesting that sexual selection may even lead to decreased cognitive ability via trade-offs with expensive secondary sexual traits.

These comparative studies provide a less than convincing case for an important role of the mating system in the evolution of cognition. Given the many, potentially confounding, factors associated with animal sociality, mating systems and ecology, it may be that controlled experiments are needed to compliment comparative methods^23^ and evaluate the generality of the mating mind hypothesis as applied to animals in general. The question thus remains whether sexual selection and mating system variation generally are important for the evolution of cognition in its widest definition – i.e. does selection for cognitive performance during mating competition lead to greater cognition when applied to other tasks or behaviors?

The mating mind hypothesis is routed in general sexual selection theory^17,24^, which holds that competition over access to mating partners should select for males that carry genomes free of deleterious mutations (i.e. males carrying the “*good genes*”)^25^. Since most new mutations are thought to be deleterious and have wide ranging pleiotropic effects on fitness related traits^26^, it follows that males that are successful in mating competition should on average be superior performers, and pass on these “good genes” to both sons and daughters^27^. Hence, the mating mind and good genes hypothesis make largely parallel predictions of an association between the mating system and general cognitive ability.

Here we tested the role of sexual selection and mating system variation in the evolution of cognition by harnessing the empirical potential of experimental evolution and the seed beetle study system. We evolved replicate evolution lines of *Callosobruchus maculatus* beetles under enforced monogamy (excluding sexual selection) or natural polygamy (including high levels of sexual competition and mate choice) for 35 generations. We then subjected these lines to a cognitively challenging spatial and chemo-sensory task, composed of mate finding and discrimination in males, and host seed finding and discrimination in females. We also assessed cognitive learning for both tasks (in males and females respectively) by letting the same beetles perform the task in three consecutive rounds with interspaced acclimation periods. Hence, our design allowed us to assess whether experimental evolution had led to improved cognitive performance in the specific task (male mate search and discrimination) known to be under differential selection in the monogamy and polygamy regime, and then to explore whether evolution under the contrasting mating systems had led to genetic changes in general cognitive ability inferred from i) improved female host search and discrimination, and ii) improved cognitive learning in the tasks, assessed in respective sex.

## Methods

### Study species

*C. maculatus* seed beetles are common pests of legumes (Fabaceae) in Africa and Asia. Females lay eggs on seeds and larvae burrow into the seed where the entire development occurs^28^. Beetles emerging from seeds are reproductively mature and require neither water nor food to reproduce (e.g.^28,29^). Adults typically die in 7-14 days after emergence in the absence of food or water (e.g.^30^).

Sexual selection is intense in this species, including both pre- and post-copulatory processes^31–36^. Sexual selection is thus likely to put demands on both male and female cognitive abilities associated with mate choice, including assessing sex, age, body size, phenotypic quality, as well as mating status of potential mating partners, as all these choices can potentially influence reproductive success ^28,31,37–39^. In the lab environment males search for females among beans, putting additional demands on male spatial orientation and use of olfactory cues to locate and discriminate females^40^, who when mated often hide amongst the beans to escape costly re-mating attempts by males^34–36^.

Female host plant search and discrimination is complex. *C. maculatus* has a wide repertoire of fabaceus host plants^41^ but females have a clear host hierarchy and preference, and they discriminate between high quality and low quality species (e.g. ^42,43^). In the lab environment females are typically presented with beans from one host species, but need to discriminate among good and bad quality seeds as well as against egg-laden seeds, as high larval density or poor quality host seeds limit both survival and size at maturity of offspring, and thus come at substantial fitness costs^38,43^.

The experimental evolution lines (see below) come from a genetic stock recently isolated from the wild^44^. *C. maculatus* utilizes both natural habitats, where host plant patches used for both egg laying and adult nectar feeding are more widely distributed, as well as grain storages, where adult food is absent but egg laying substrate and population density is higher^38,41,43^. Thus, selection on spatial cognition and chemosensory cues is likely always strong in natural populations, but may take different forms, which is predicted to maintain genetic variation in the cognitive traits under study, as seen for other characters related to these alternative environments (e.g.^45^).

### Polygamous and Monogamous experimental evolution lines

The lines used in this study are thoroughly described in Martinossi-Allibert et al. ^46^. In brief, the lines and the outbred base population from which they originate were maintained under controlled temperature (29°C), humidity (50%RH) and light cycle (12L: 12D), and reared on the preferred host plant^41^ *Vigna unguiculata* (black-eyed bean). We used six lines in this study; three replicate lines evolving under enforced monogamy (removing sexual selection), and three lines evolving under polygamy in the natural lab environment which applies sexual selection by allowing pre- and post-copulatory mate competition and choice, in addition to the fecundity and viability selection acting in the monogamy regime. There are two more lines in the study by Martinossi-Allibert et al. from a third evolution regime which applied sexual selection on males while excluding selection on adult females all together. However, since our hypotheses were most straightforward to test by comparing the effect of adding sexual selection (polygamy regime) to an already natural socio-sexual mating system (monogamy regime), we did not include these two lines in this study.

Effective population size in each regime was kept approximately equal (N_e_ = 150, N_monogamy_ = 246, N_polygamy_ = 300) and the number of beans provided as egg laying substrate in each regime was standardized to give the same, relatively low, juvenile density (2-4 eggs/bean) to minimize (and equalize) larval competition^46^. The polygamy and monogamy regime show differences in line with good genes effects of sexual selection: already following 16-20 generations of experimental evolution, polygamy lines showed higher lifetime reproductive success as well as population fitness compared to monogamy lines^46^.

### Cognitive performance of males in the mate searching task

We measured focal males’ (i.e. derived from one of the evolution lines) ability to localise and discriminate females in a spatially complex arena made up by a petri dish measuring 150 mm in diameter containing reference beetles of both sexes (Fig. 1). The reference beetles originated from the original base population from which the selection lines were derived. Virgin reference beetles, 0-24h old, were frozen at −20°C and defrosted just previous to a trail and glued to the arena (with their ventral side facing downwards), making sure that the focal beetles did not shift their position during the three consecutive behavioural trails. The same reference beetles were used for the full run of three trails before being replaced by new beetles.

**Figure 1:**
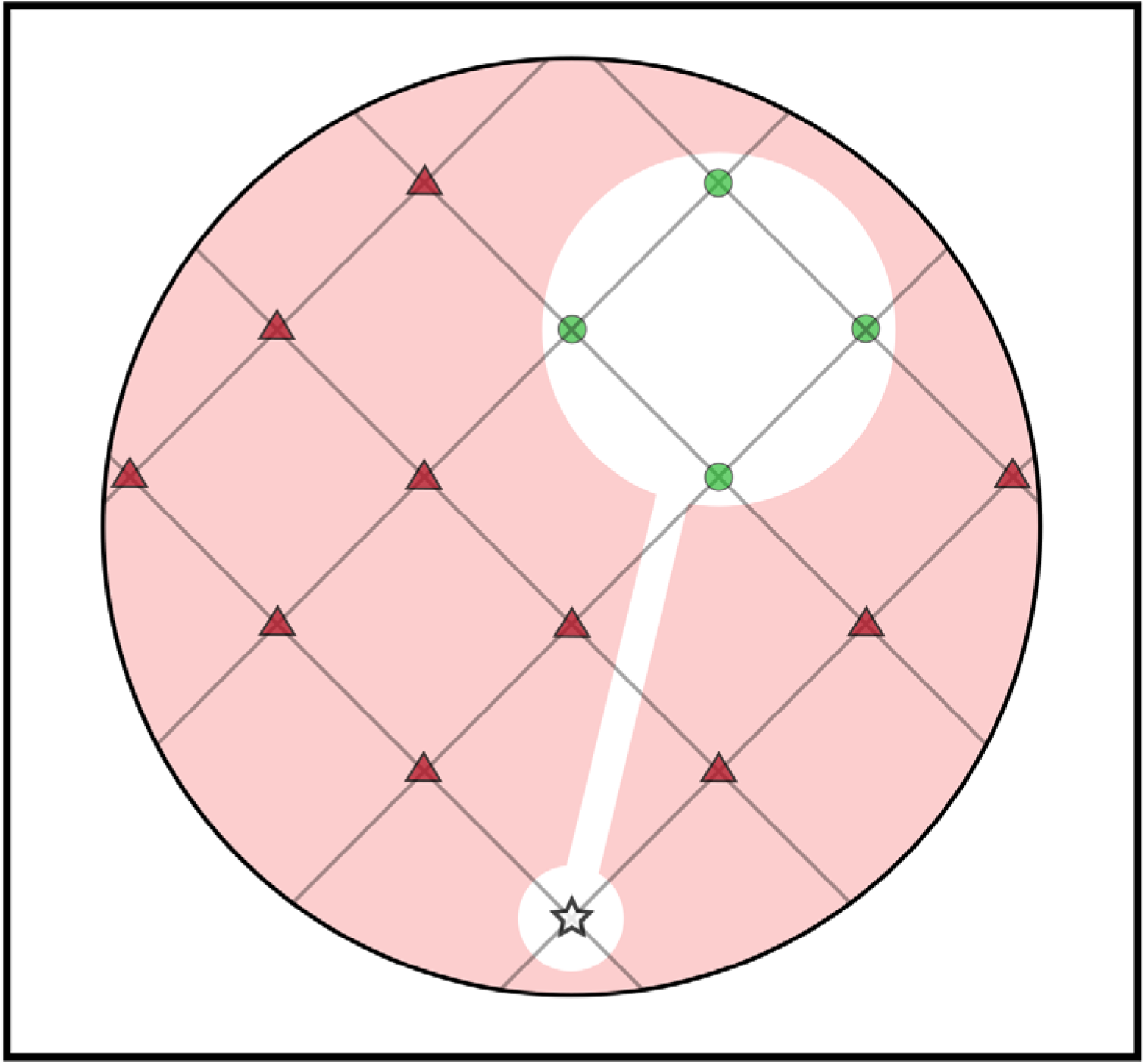
The experimental arena (a 150 mm diameter petri-dish) used for the behavioural assays. The symbols indicate the following: white star = place where the four focal individuals were placed at the start of each trail; red triangles = the wrong choice (males for male trails | chick-peas for female trails), green circles = the correct choice (females for male trails | black-eyed beans for female trails).

The arena floor was covered with paper (Fig. S1) designed to aid spatial memory and learning. The paper had pale red background and a white large circle connected to a white channel leading away from it towards the inner wall of the opposite side of the arena, ending in a smaller circle at which the four focal males were placed at the initiation of each trial (see below). Each large white circle contained four equidistant points where the freshly defrosted reference females were glued. The ‘channel’ was connected to the circle to easier allow focal males to learn and find the location of females. The remaining pale red background comprised ten equidistant points where the freshly defrosted reference males were glued (Fig. 1). A second type of arena with inversed color scheme was used to control for effects of potential color preferences in the beetles (Fig. S1).

The assays were run on a heating plate set at 30°C, with six arenas on the plate scored simultaneously. In each arena, 4 focal males were introduced simultaneously. This was for two reasons, i) to increase activity, because beetles become more active in group, and ii) to efficiently score as many beetles as possible. Assays were initiated by consecutively introducing the four beetles into the small circle in each of the 6 replicate arenas. This took ca. 60 seconds after which behavioral observations were taken in the same sequential order each minute for the subsequent 10 minutes. Each census time of an arena lasted for 10 seconds before the next arena in line was observed. During the 10 seconds we recorded whether each of the four beetles in the arena were in contact with a reference individual and whether this was a male or a female. This contact usually meant that males were trying to, or even “successfully” mated with both (dead) males and females (males try to mate readily with other males in this species and population^40^). The four beetles were thus scored as a group and could at each census time get a score between 0 and 4 for mating attempts on reference males and females.

Our measurements are likely to capture two independent aspects of seed beetle cognition; i) the ability to locate and remember the location of reference individuals in a spatially complex two-dimensional landscape, and ii) via chemo-sensory cues discriminate the sex of the located reference individual (which seems cognitively demanding in seed beetles:^40^, as in many other insects^47^).

### Cognitive performance of females in the host searching task

We measured focal females’ ability to find and discriminate among a high quality (*V. unguiculata*, black-eyed bean) and a low quality (*Cicer arietinum*: Chick pea) host species^41^ in the exact same type of arena and set-up (Fig. 1, Fig. S1). To make sure that females were motivated to search for hosts, they were mated with conspecific males 24 hours prior to the trials, and prevented from laying eggs by depriving them of host seeds. In contrast to male trails, host seeds did not need to be glued to stay in their place. Host seeds were also removed between each female trail as eggs were laid readily by females and the presence of eggs on the hosts can affect female egg laying behaviour (heavily laden seeds are more often rejected). We registered female host inspection as behaviour. This inspection behavior was usually in form of females being on top of seeds making tactile contact with or ovipositing on the seed. Females readily laid eggs on both types of hosts during the assays, although this was never quantified. Similarly to the male assays, our scoring of behaviour captures variation in female cognition in terms of i) spatial orientation and memory as well as ii) chemo-sensory cues associated with host discrimination.

### Scoring behavior

The observer (JdA) was always blinded to which line and evolution regime that was assayed. Each group of four beetles were scored for their behaviour as a group through three 10 minute trails with a 20 minute acclimation period in a 30mm diameter petri dish before and in-between each trail. We could thus study overall differences in beetle cognition in terms of the average performance over time in each line and sex, as well as the potential for cognitive learning by looking at the improvement in performance within and between trails. We used two heating plates, each with one of the two color schemes (Fig. S1). The heating plates were run interchangeably (during the other plate’s acclimation periods), and each line was scored once on each plate/color scheme per sex, resulting in 12 arena replicates per sex and line. Thus, 48 males and 48 females from each of the 6 lines were scored for behaviour over three consecutive 10 minute trails for a total of 17.280 behavioral observations.

### Statistics

Male and female data were of the same form and were first analysed separately in equivalent models. We modelled the response as an “error rate” (the fraction of incorrect choices) over the trails using a binomially distributed response variable with the levels “male/female” (male assays) or “good host/bad host” (female assays). Time of the trail was included as a covariate that was linearized by taking its natural logarithm prior to analysis. Evolution regime and trail number were analysed as discrete factors. We included interactions among all three of these explanatory variables, where two or three-way interactions including evolution regime would signify differences in performance over time among the two evolution regimes, indicative of an effect of sexual selection on cognitive learning. A main effect of evolution regime, on the other hand, would indicate an effect of sexual selection on general cognitive performance in the given task. Line identity was included as a random effect crossed with the three explanatory variables to account for the true replication of the experiment (being the six line replicates, and not individual observations). We included assay identity (the four beetles run together over consecutive trails) as an additional random effect. We also modelled main effects of heating plate to control for spatial effects in the lab and beetle color preferences. However, this effect was never significant and was ultimately removed from all models.

In addition to the male- and female-specific model, we also looked more formally for sex-differences in the response of cognitive performance to experimental evolution under alternative mating systems. This was done by recoding the levels of the response variable to “correct” and incorrect” and then running a statistical model with the main effect of “sex” crossed by the three other explanatory variables of interest (time, trail number and evolution regime) and the random effect of line identity.

Analyses were carried out with the package lme4^48^ in the statistical software R. We used the “bobyqa” optimizer to increase the number of iterations (to 100.000) to achieve convergence of all models. We report type-II P-values based on likelihood ratio tests and χ^2^-statistics.

## Results

### Cognitive performance of males

Throughout the time of the trials, males increased the fraction of mounting attempts on females (χ^2^_1_= 80.1, P < 0.001), suggesting that they learned the spatial location of females and then preferred to stay there to try to mate with them. While there was no main effect of trial, there was a strong interaction between trail and time (χ^2^_2_= 18.7, P < 0.001) because males were more efficient in finding females already at the start of the third trail when trained, compared to the first trail when naïve. These results thus demonstrate clear effects of learning on performance in the mate searching task (Fig. 2A & C, Fig. S2). Males from the polygamy regime were more efficient overall in discriminating the sex of beetles compared to males from the monogamy regime (χ^2^_1_= 8.56, P = 0.003, Fig. 2A & C). This difference was mainly driven by the number of mounting attempts on reference males, which was much higher in males from monogamous lines (Fig. S2). However, there were no significant differences between selection regimes in how mate discrimination changed within or between trails (Fig. 2A & C, Table S2, Fig S2), indicating that learning was similar in the two evolution regimes (full statistics in Supplementary Table S2).

**Figure 2:**
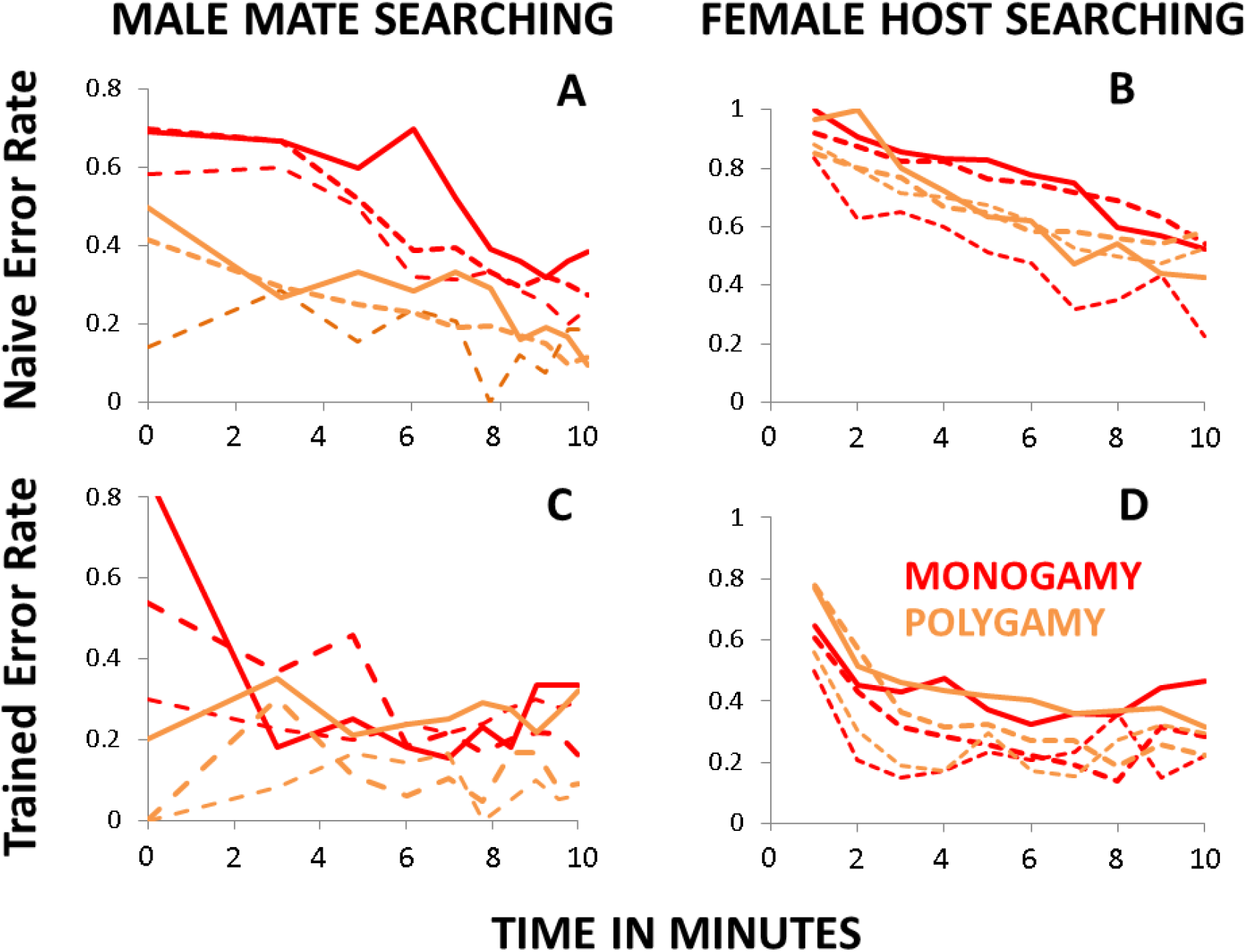
Sex-specific baseline cognitive performance and learning in the three replicate monogamous (red) and polygamous (orange) lines. Male mate searching ability (A, C) and female host searching ability (B, D) in terms of “error rates” (the fraction of male mating attempts with other males, and the fraction of female contacts with the suboptimal host). Shown are data for naïve beetles (A, B) in the first 10-minute trail, and trained beetles (C, D) in the third 10-minute trail. The line type (thick and full to thin and hatched) designates the line identity and makes it possible to match line performance across the first and the third trail.

### Cognitive performance of females

Throughout the time of the trials, females decreased the fraction of visits to suboptimal host seeds (χ^2^_1_= 246.7, P < 0.001), suggesting that they learned the spatial location of the optimal host and then preferred to stay there to oviposit (Fig. 2B, D, Fig. S3). There was also a strong main effect of trial (χ^2^_2_= 99.1, P < 0.001), as well as a strong interaction between trail and time (χ^2^_2_= 33.8, P < 0.001), signifying that females improved in the host searching task through spatial and/or chemo-sensory learning (Fig. 2B, D, Fig. S3). However, contrary to the superior performance of polygamous lines in the male task, females from the two regimes did not show any overall differences in host search and discrimination (Table S3). This sex-difference in the evolutionary response of cognitive performance was statistically significant (χ^2^_1_= 5.91, P = 0.015, Table S4). There were also no significant two-way interactions between selection regime and time or trail (Table S3), suggesting that female learning was largely similar in the two evolution regimes. There was, however, a marginally nonsignificant three-way interaction between selection regime, trail and time (χ^2^_2_= 5.76, P < 0.056, Fig S3). This trend was driven by a pattern where evolution regimes showed very similar performance throughout the first trail as naïve beetles (regime:time interaction: χ^2^_1_= 0.02, P = 0.89), while monogamous lines tended to be better at discriminating between hosts when trained at the start of the third trail, but where this difference between regimes quickly disappeared as the trail went along (regime:time interaction: χ^2^_1_= 4.28, P = 0.039, Fig. S3). We note that this difference runs counter to the expectation that sexual selection should improve general cognitive abilities (full statistics in Supplementary S3 & S4).

### Sex-specifíc correlations between cognitive performance and lifetime reproductive success

We explored the link between cognitive performance and fitness in each sex by calculating correlations between lifetime reproductive success (LRS, reported in^46^) and the measured cognitive traits (error rate in trail 1-3 and a learning score based on the relative reduction in error rate between the first and last trail: [e1-e3] / e1), based on trait means per sex and evolution line replicate. These correlations are graphically depicted in figure 4. We note that our interpretation here must remain tentative since these correlations are based only on 6 data-points (three replicate lines per mating regime), and hence, cannot be used for rigorous statistical testing. Error rates are highly correlated between the three trails within each sex (r = 0.58-0.96), suggesting substantial repeatability in behaviour among the six genotypes. However, error rates and the learning score are very weakly correlated between sexes (r = −0.08−0.34), implying that different genes govern mate search in males and host search in females. Moreover, while male LRS was negatively correlated to male error rates (r = −0.68 − −0.59) and positively correlated to male learning (r = 0.59) as predicted, it was positively correlated to the female error rate (r = 0.30-0.62) and negatively correlated to female learning (r = −0.51). This may suggest genetic conflict between the sexes at loci encoding the studied cognitive traits. Finally, female learning was negatively correlated to female LRS on both the focal ancestral host (black eyed bean; r = −0.68) as well as an alternative host^46^ (adzuki bean; r = −0.71), suggesting that cognitive learning may trade-off against fecundity in females.

## Discussion

It is well known that sexual selection can put demands on cognitive abilities related to sexual signalling and mate choice^5,16–18^ and mate search has also been linked to spatial learning in both vertebrates^49^ and insects^50^. For example, in guppies, females from lines selected for larger brains were better at choosing among high and low quality mating partners^51^ and males from the same lines were better at finding mates in a spatial learning task^52^. Similarly, in fruit flies, cognitive learning improves female mate choice^18^. However, whether the mating system can drive species differences in general cognition^13^ is much more disputed and direct evidence remains scarce. Here we have shown that mating system variation can lead to the evolution of cognitive performance. Males evolving under polygamy were more efficient in directing their mating effort towards females in spatially complex mixed-sex settings. Given that these males also have higher reproductive success than males evolving under monogamy^46^, this suggests that increased cognitive performance in mate search has fitness benefits in males (see also Fig. 4). However, the evolved increase in mate search ability was not accompanied by improved learning or increases in female cognitive performance, as expected under the mating mind hypothesis.

These negative results are readily interpretable as there was sufficient power in our design to detect significant differences in male performance between evolution regimes, as well as to demonstrate substantial improvement in the cognitive tasks through learning in both males and females (Figs. 2 & 3). Our main results thus imply that good genes processes resulting in overall improvement of cognitive ability may not materialize when sexual selection acts on standing genetic variation, as it did in our experiment. Interestingly, the decreased ability of monogamous males (relative to polygamous males) to avoid directing mating attempts toward other males was drastic and evolved in only 35 generations of relaxed sexual selection (Fig. S2). This fast decrease implies that selection acted on segregating genetic variation with antagonistic pleiotropic effects on other fitness related traits, because i) decreases in monogamous lines due to the accumulation of de novo mutation over such short time frames seem unlikely, and ii) effects of genetic drift should be negligible since effective population size was relatively large (Ne ~150), all three replicate lines for each regime showed parallel divergence (Fig 2A & C, Fig. 3) and the effect of drift was controlled for in the applied statistical models. Our results thus also pose the question of what maintains such vast amounts of genetic variation in male cognitive behaviour.

**Figure 3:**
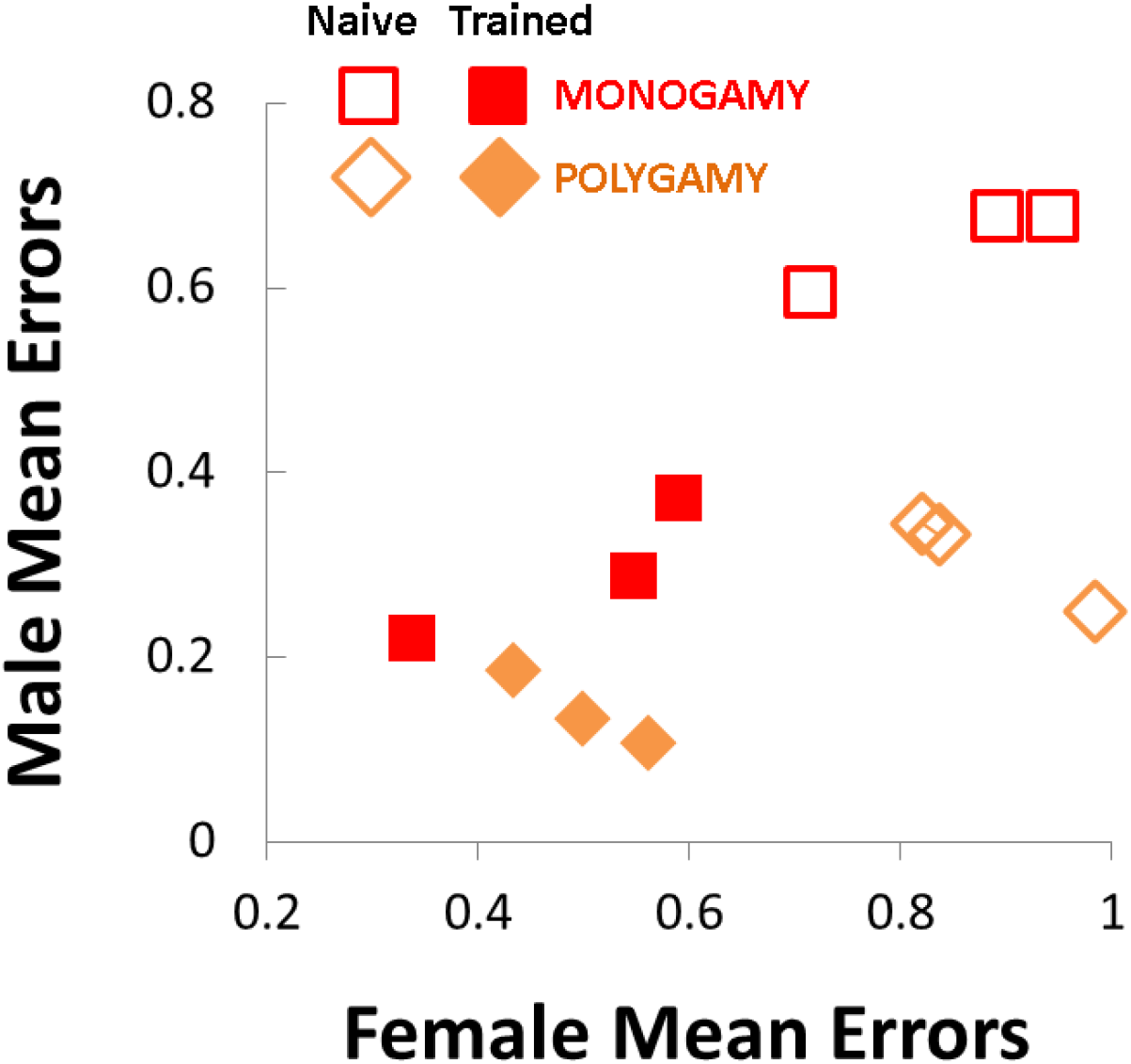
Cognitive performance (in terms of error rates) averaged over the full 10-minute trial, for naïve beetles in the first trail (open symbols) and trained beetles in the third trail (closed symbols). Mean male and female error rates are shown for each of the three replicate monogamous (red) and polygamous (orange) lines.

Our results are in many ways similar to the study by Hollis and Kawecki (2014) on *Drosophila melanogaster* fruit flies, which is the only other study we know of that has applied experimental evolution and manipulated the mating system to look at effects on cognitive abilities. In their study, evolution under polygamy contributed to the maintenance of mate acquisition abilities in males, but also lead to superior aversive learning - a cognitive task not directly related to the applied sexual selection. While this suggests that sexual selection improved general cognitive abilities, in line with the mating mind hypothesis, polygamous females showed no such increase, and even tendencies for reduced cognitive performance relative to females from monogamous lines^53^. This is also in line with our results, showing no differences in performance between monogamous and polygamous *C. maculatus* females overall, and a tendency for monogamous females to learn faster (Fig. S3).

Indeed, as an alternative to good genes effects, sexual selection may lead to sex-limited responses and increased sexual dimorphism in cognition^13,19,49,54,55^. Such an outcome is expected when males and females experience different selection pressures and genetic constraints are not insurmountable^56^, so that cognitive traits can evolve independently in each sex^19^. One mechanistic explanation for the sex-specificity observed in this study could be differences in the chemosensory machinery required to successfully identify and discriminate the sex of mating partners and host species (in males and females respectively). Given that collecting and processing such information should require costly development and maintenance of neuroreceptors^4,50^, cognitive performance in mate search and host search may trade-off against each other, if different receptors are employed for the two tasks and these receptors compete for resources, physical space, or downstream cognitive processing of their transmitted information. This hypothesis is in line with the tendency for sexual selection to have positive effects on male mate search but slightly negative effects on female cognitive learning in both our study and Hollis and Kawecki’s (2014) study on fruit flies. Moreover, in this population of beetle, female fecundity has previously been shown to be negatively correlated to the accuracy of male sex discrimination^40^, and in this study, female host search and discrimination tended to be negatively correlated to male reproductive success (Fig. 4).

**Figure 4:**
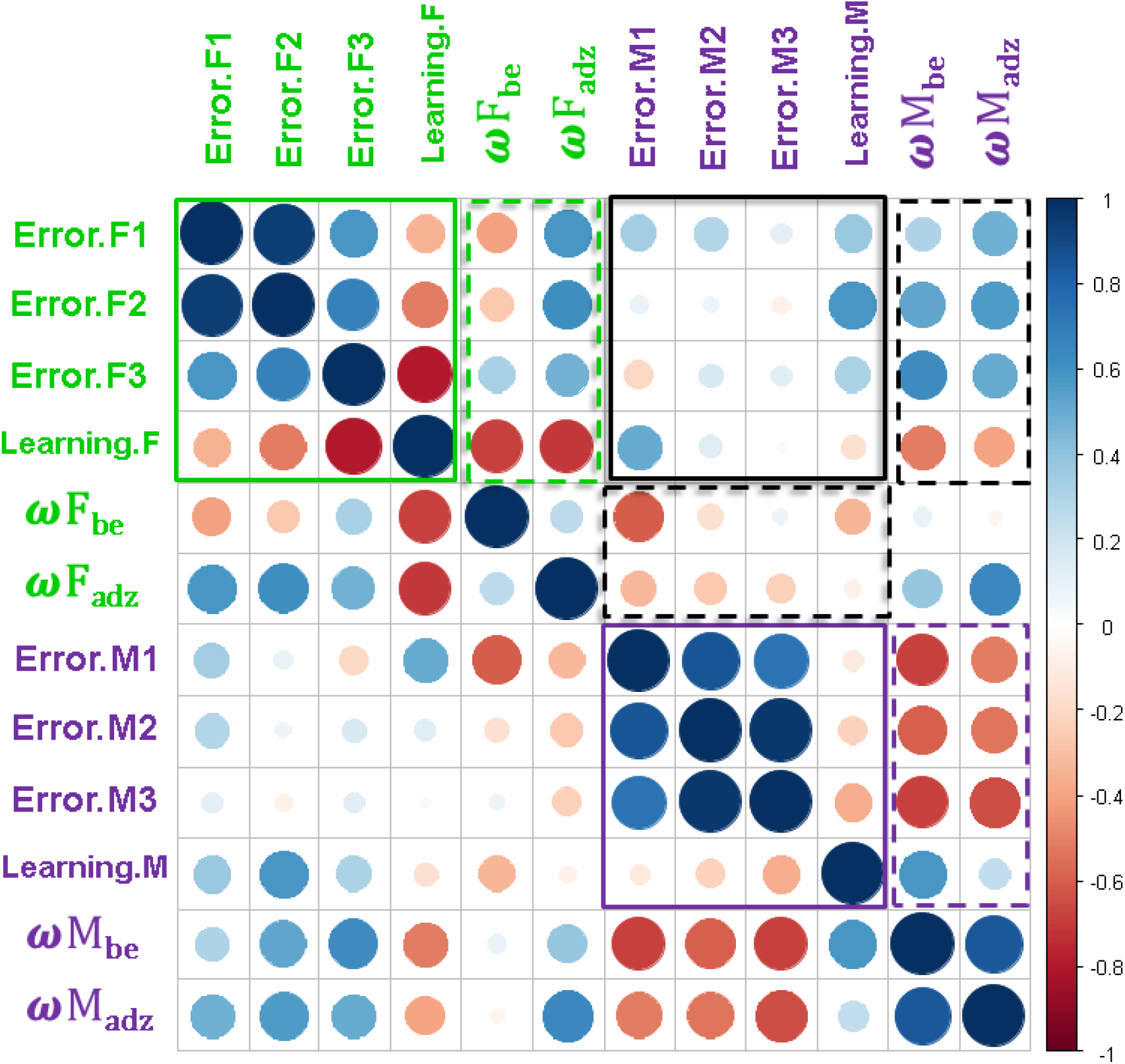
Sex-specific genetic correlations between cognitive performance and lifetime reproductive success. Shown are correlations based on the 6 lines (3 from each regime) between cognitive performance (error rates in the three consecutive behavioral trails and a measure of learning from trail 1 to 3: [E1-E3]/E1), and lifetime reproductive success () on two host species (be = ancestral black eyed beans, adz = adzuki beans). Within-sex genetic correlations are highlighted by green (F = female) and purple (M = male) squares. Black squares highlight between-sex genetic correlations. Full lines designate correlations between measures of cognitive performance and hatched lines between cognitive performance and lifetime reproductive success. Note that the same correlations are depicted both above and below the diagonal. Circles on the diagonal are trait variances standardized to a size = 1.

These results imply that selection on cognitive traits may sometimes act with opposing forces in the sexes. If in such cases genetic constraints are preventing each sex from evolving independently from the other^56,57^, this type of sexual antagonism^58^ will generate balancing selection that can act to maintain allelic variation at genes underpinning cognitive abilities^59,60^. Mechanistically, sexual antagonism over cognitive traits could, for example, arise if males benefit mostly from increasing allocation to one type of chemoreceptor (e.g. increasing accuracy of sex discrimination) while females benefit from allocation to another type of receptor (e.g. increasing accuracy of host discrimination). Sexual antagonism could also arise if the benefit of a specific cognitive ability is limited mainly to one sex while its energetic cost is paid by both sexes, as seen for other types of traits under sexually antagonistic selection^40,61,62^. In this study, female learning correlated negatively with both male and female reproductive success (Fig. 4), in line with this hypothesis.

Indeed, cognitive traits and learning are generally assumed to come with energetic costs. For example, there are cost of developing and using a large brain in vertebrates^7,9,63^, as well as documented costs of memory and allocation to cognitive traits in insects^4,50,64–66^. Similarly, sexually selected traits are themselves expected to be costly ^17,24,27,67^, and while some studies have found a positive genetic correlation between primary and secondary sexual traits and brain size (e.g.^68,69^), in line with good genes effect, there are also examples of negative correlations (e.g.^22^), suggesting that increased sexual selection may sometimes lead to decreases in cognitive traits via energy allocation trade-offs^24,70,71^. The action of such antagonistic pleiotropy within and between sexes to maintain genetic variation could thus be responsible for the substantial amounts of standing genetic variation in male sex discrimination documented here and previously^40^ in this population of *C. maculatus*. The notion that sexually antagonistic selection has played a key role in this process is also supported by previous studies on the population^40,44,72,73^.

Behavioral plasticity can play an important role in deciding species distributions, persistence and modes of adaptation to changing environments, for example by increasing the efficacy of spatial exploration and resource sampling mediating niche matching^1–5^. Cognitive processes are also key in mate choice dynamics and may therefore play a role in speciation^74–79^. The interplay between sexual selection and the evolution of cognition, with special emphasis on potentially underappreciated effects of sexually antagonistic selection on cognitive traits, may therefore have important consequences for evolutionary dynamics and certainly deserves more attention in other study systems. While the mating mind and good genes hypothesis predict a positive association between cognitive ability and the strength of sexual selection to be built up by purifying selection against recurrent deleterious pleiotropic mutations, our study implies that much of the standing genetic variation in cognitive performance upon which evolutionary responses to novel environments rely, will have been moulded and maintained by forces of balancing selection within and between the sexes. This sets the stage for rapid sex-specific responses to changes in ecological and socio-sexual conditions.

### Ethics statement

All experiments have been carried out within the regulations held by Swedish governmental laws. Actions were taken to reduce the number of beetles used in the study.

### Data accessibility

Data will be uploaded to the Dryad Data Repository upon potential acceptance

### Author’s contributions

The study was conceived by JB, JdA and DB, and was carried out by JdA and JB. DB analyzed the data and wrote the manuscript. All authors commented on the first draft.

## Acknowledgements

We thank Ivain Martinossi-Allibert and Göran Arnqvist who have helped to create the experimental evolution lines. We also like to thank Johanna Liljestrand-Rönn for practical help in the lab.

## Supplementary Material

**Figure S1:**
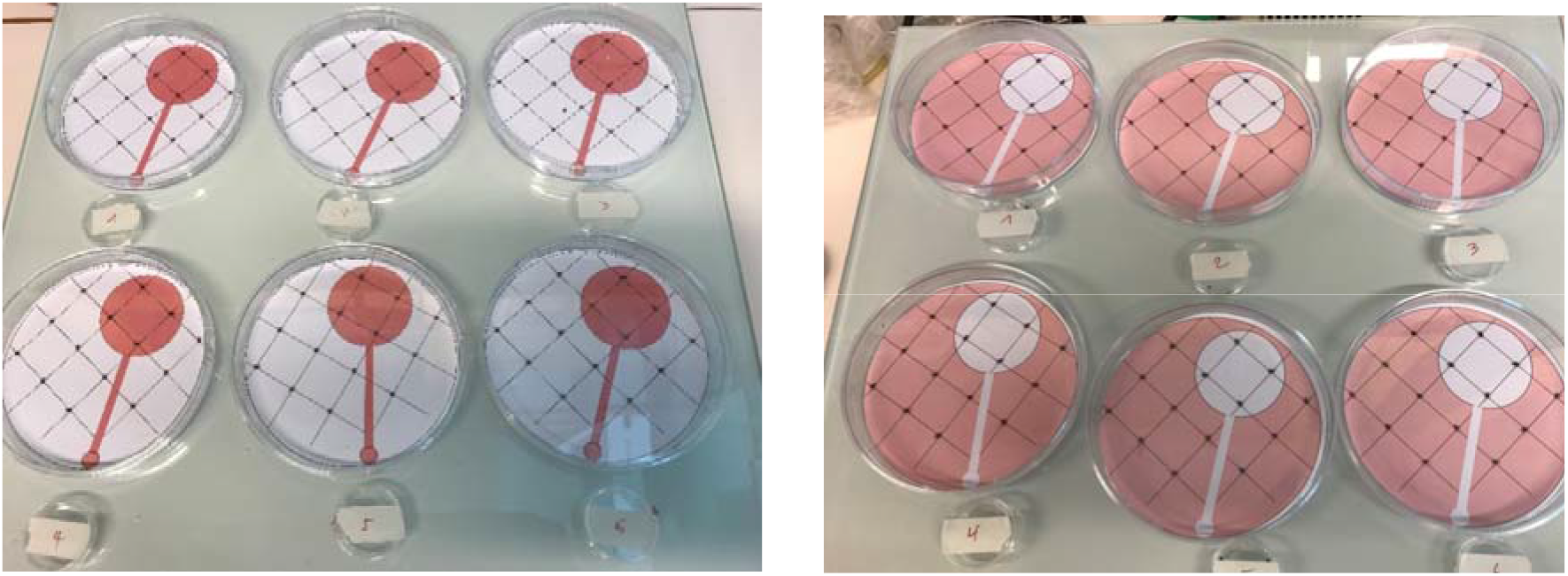
Experimental arenas. Arenas used to assess cognition and learning. Shown are the two arena types with reversed color schemes, placed on heating plates situated ca. 1 metre apart. Below the arenas are the 30mm diameter acclimation petri-dishes where the four focal beetles spent 20 minutes prior to and in-between trails.

**Figure S2:**
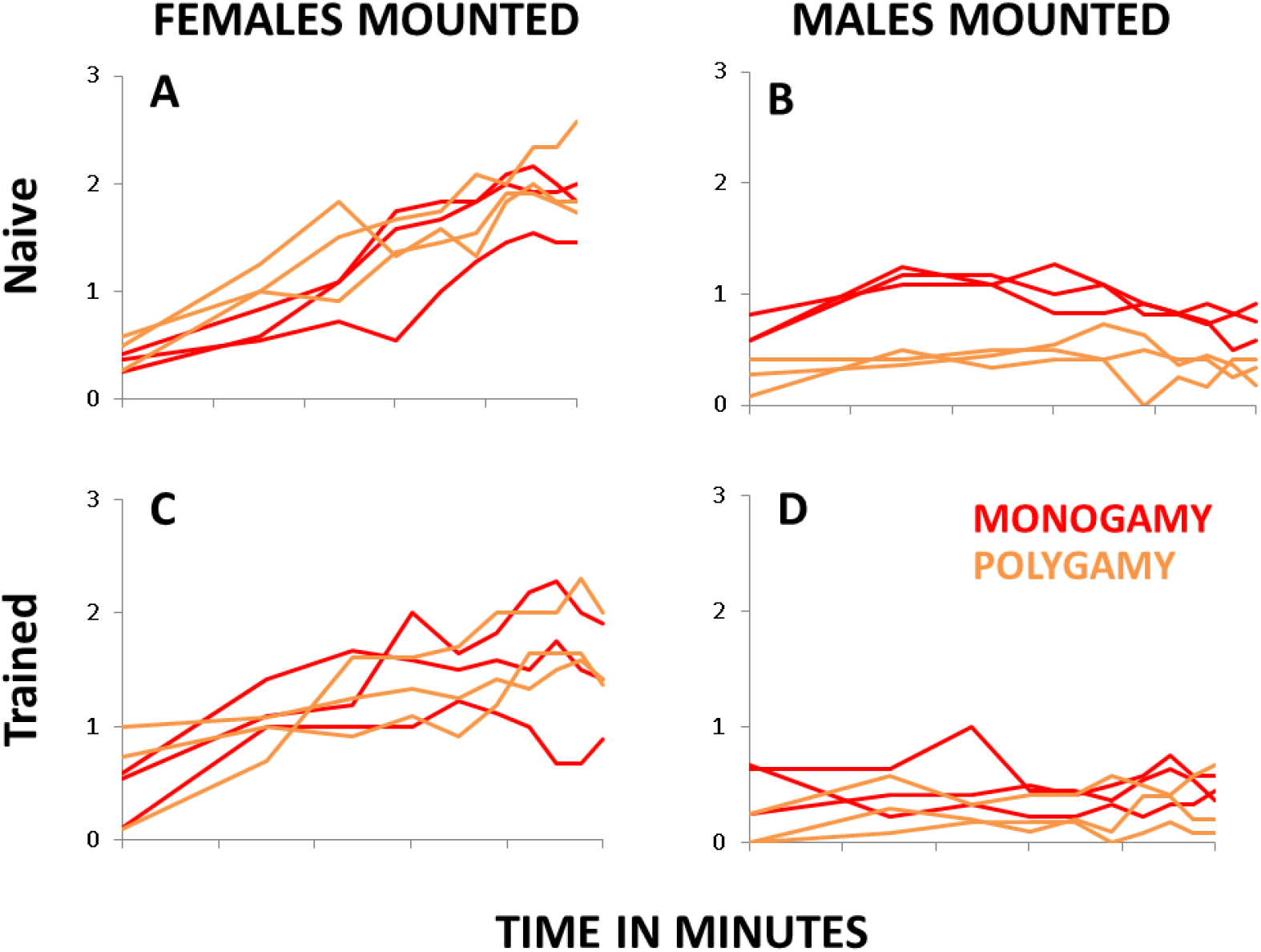
Male cognitive performance. The averaged summed number of times that the four males made mounting attempts on females (left; A & C) and males (right; B & D) during the first (“Naïve” beetles) and third (“Trained” beetles) behavioural trail.

**Table S2:** Male cognitive performance

Type II Wald chi-square tests
|  | Chisq | Df | P |  |
| --- | --- | --- | --- | --- |
| trial | 4.2328 | 2 | 0.120466 |  |
| time | 80.0619 | 1 | < 2.2e-16 | *** |
| regime | 8.5645 | 1 | 0.003428 | ** |
| trial:time | 18.7180 | 2 | 8.619e-05 | *** |
| trial:regime | 3.8489 | 2 | 0.145959 |  |
| time:regime | 2.3646 | 1 | 0.124115 |  |
| trial:time:regime | 0.1789 | 2 | 0.914416 |  |

**Figure S3:**
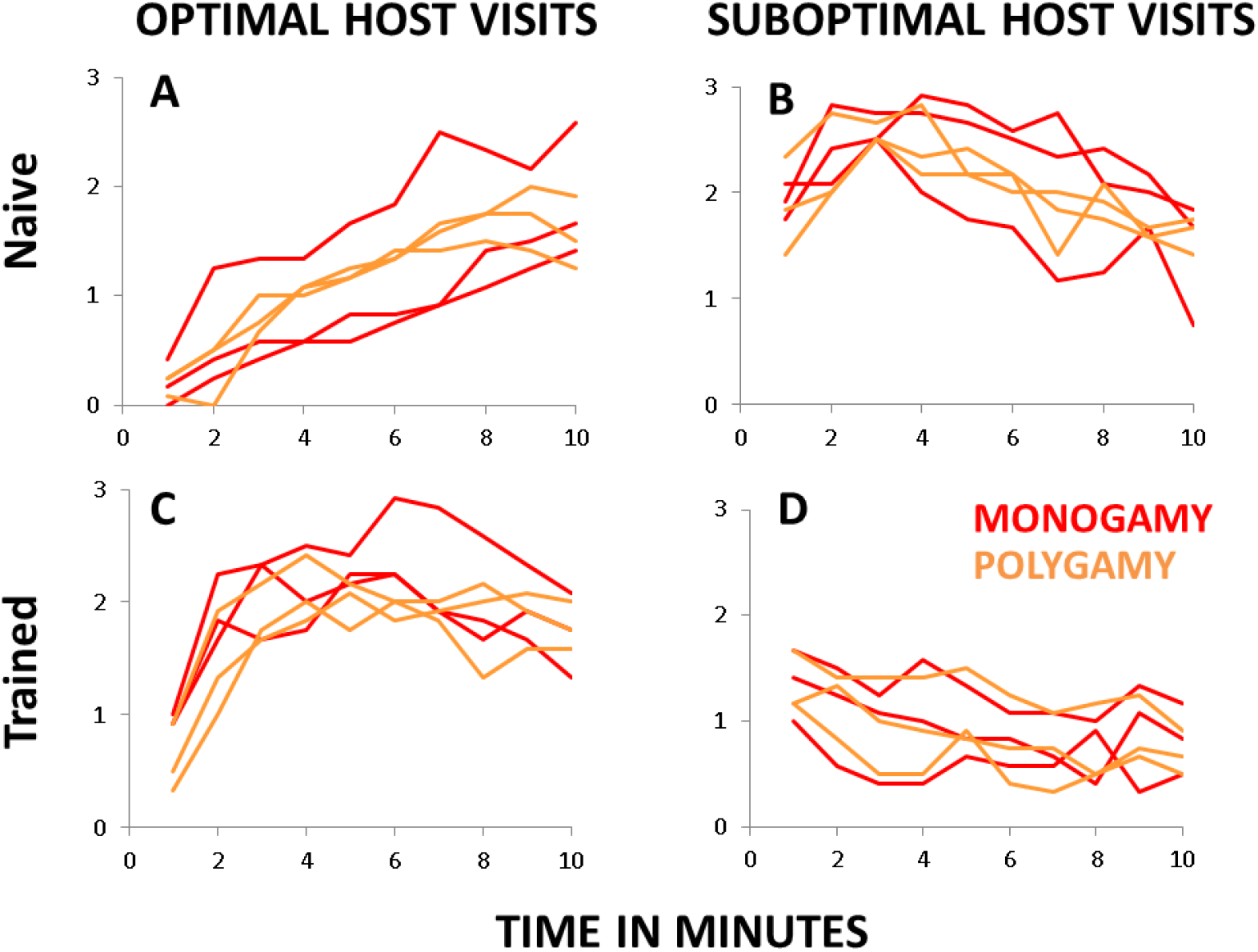
Female cognitive performance. The averaged summed number of times that the four females made inspections of optimal black-eyed beans (left; A & C) and sub-optimal chick-peas (right; B & D) during the first (“Naïve” beetles) and third (“Trained” beetles).

**Table S3:**
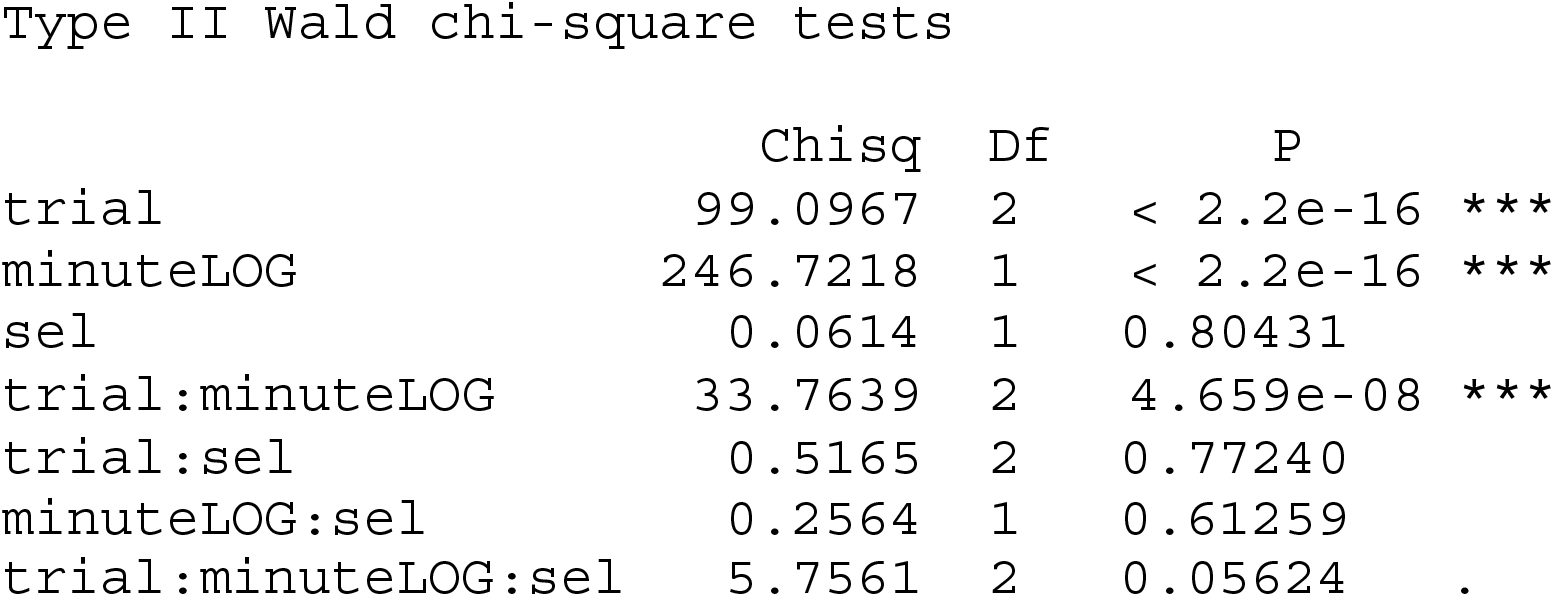
Female cognitive performance

Type II Wald chi-square tests
|  | Chisq | Df | P |  |
| --- | --- | --- | --- | --- |
| trial | 99.0967 | 2 | < 2.2e-16 | *** |
| minuteLOG | 246.7218 | 1 | < 2.2e-16 | *** |
| sel | 0.0614 | 1 | 0.80431 |  |
| trial:minuteLOG | 33.7639 | 2 | 4.659e-08 | *** |
| trial:sel | 0.5165 | 2 | 0.77240 |  |
| minuteLOG:sel | 0.2564 | 1 | 0.61259 |  |
| trial:minuteLOG:sel | 5.7561 | 2 | 0.05624 | . |

**Table S4:**
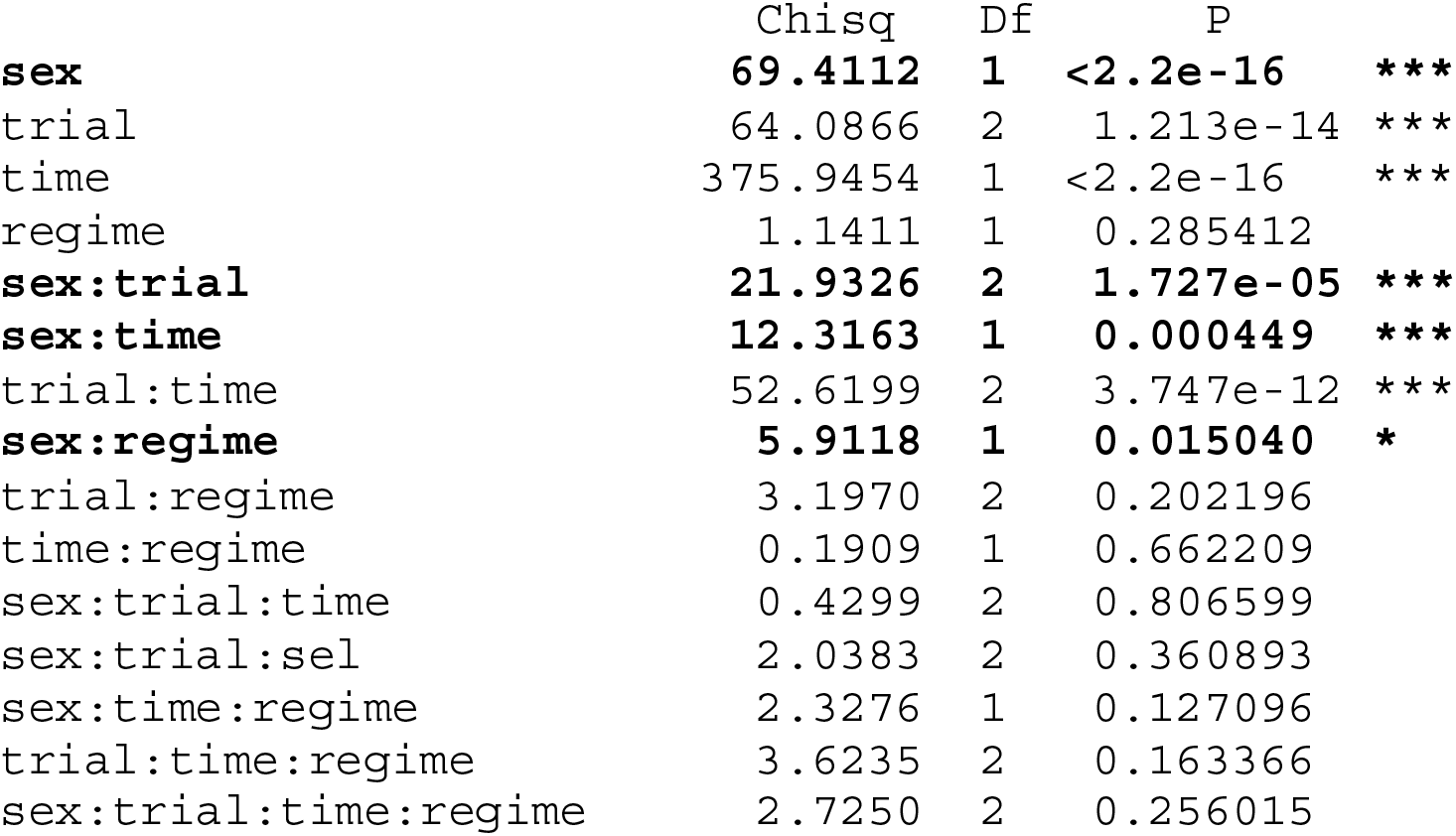
Sex differences in cognitive performance

